# Systematical analysis reveals the novel function of *Cyp2c29* in liver injury

**DOI:** 10.1101/763581

**Authors:** Qi Wang, Qin Tang, Lijun Zhao, Qiong Zhang, Yuxin Wu, Hui Hu, Lan-Lan Liu, Xiang Liu, Yanhong Zhu, An-Yuan Guo, Xiangliang Yang

## Abstract

As a severe lethal cancer, hepatocellular carcinoma (HCC) usually originates from chronic liver injury and inflammation, in which the discovery of key genes is important for HCC prevention. Here, we analyzed the time serial (from 0 week to 30 weeks) transcriptome data of liver injury samples in diethylnitrosamine (DEN)-induced HCC mouse model. Through expression and function analyses, we identified that *Cyp2c29* was a key gene continuously downregulated during liver injury. Overexpression of Cyp2c29 suppressed the NF-κB activation, proinflammatory cytokine production and hepatocyte proliferation by increasing its production 14,15-epoxyeicosatrienoic acid (14,15-EET). Furthermore, *in vivo Cyp2c29* protected against liver inflammation in liver injury mice models by reversing the expression on functions of cell proliferation, metabolism and inflammation including suppressing NF-κB pathway and compensatory proliferation. *CYP2C8* and *CYP2C9*, two human homologs of mouse *Cyp2c29*, were decreased in human HCC progression and positively correlated with HCC patient survival. Therefore, through systematical analysis and verification, we identified that *Cyp2c29* is a novel gene in liver injury and inflammation, which may be a potential biomarker for HCC prevention and prognosis.

## Introduction

Hepatocellular carcinoma (HCC) is the sixth most frequently diagnosed cancer and the second most common cause of cancer-related death worldwide (Ferlay et al, et al; Colombo & Maisonneuve *et al*). Although the pathophysiology of HCC is not clearly understood, it is well known that HCC typically occurs in chronically injured and inflamed livers. Persistent injury is characterized by inflammatory cell infiltration accompanied by cytokine production (He&Karin, 2011; Balkwill&Mantovani, 2001). Chronic inflammation triggers compensatory cell proliferation and causes genetic mutations that may induce carcinogenesis (Ray, 2018). The identification of key molecules and mechanisms from the liver injury to HCC will be important for HCC prevention and diagnosis.

Many inflammation-related genes in mouse are strongly correlated with their orthologous genes in patients with liver diseases (Tacke & Zimmermann, 2014). CYP enzymes are xenobiotic-metabolizing enzymes that are primarily found in liver cells of rodents and humans (Li *et al*, 2017; Yu *et al*, 2015; Nebert & Dalton, 2006). These enzymes are involved in eicosanoid metabolism and regulate a diverse set of homeostatic and inflammatory processes. The expression levels of CYP genes are closely associated with the progression of liver diseases. For example, *Cyp2c29, Cyp2c50, Cyp2c55* and *Cyp2j5* transcription levels have been found to be suppressed in a mouse model of nonalcoholic steatohepatitis (Schuck *et al*, 2014). In addition, decreased expression of CYP genes, such as *CYP2C8, CYP2C9*, and *CYP2C19*, has been observed in HCC patients (Tsunedomi *et al*, 2005; Xu *et al*, 2001; Wang *et al*, 2018). However, the role of CYP genes in the progression of HCC is not well characterized.

High-throughput sequencing combined with bioinformatics analysis provides a systematic way to explore crucial genes and mechanisms in complex diseases (Zhou *et al*, 2018). Previously, we investigated miRNA and transcription factor (TF) coregulatory networks to reveal regulators in blood diseases (Lin *et al*, 2015; Lin *et al*, 2015; Ye *et al*, 2012). Our findings indicated that complex regulatory relationships involved in disease progression can be illuminated by transcriptome and network analysis.

In the present study, we detected differentially expressed genes (DEGs) during the progression of liver injury using time series gene expression analysis. We focused on *Cyp2c29*, which was found to be significantly downregulated in a mouse model of liver injury. Our results demonstrated that *Cyp2c29* may serve as a new biomarker for the prevention, diagnosis and prognosis of HCC.

## Results

### Transcriptome changes during the liver injury of DEN-induced mouse model

To mimic the process of hepatocarcinogenesis, low doses of DEN and high-fat diets (HFD) were used to establish a mouse model of liver injury. Liver tissues were collected at week 0, week 10, week 20, week 25, and week 30 (w0, w10, w20, w25 and w30), respectively. Mice showed body weight loss, hepatocyte necrosis and inflammatory infiltration from w10 onwards when given DEN and fed with HFD (Fig 1A, B). At w25, severe liver fibrosis was detected in the DEN treatment group, which was confirmed by observation of extensive collagen deposition and bridges between vessels upon Masson trichrome staining (Fig 1C). At w30, tumors were observed in 5 out of 6 mice in the DEN treatment group. In addition, hematoxylin and eosin (H&E) staining of tumor tissues showed a typical trabecular growth pattern (Fig 1B).

**Figure 1.**
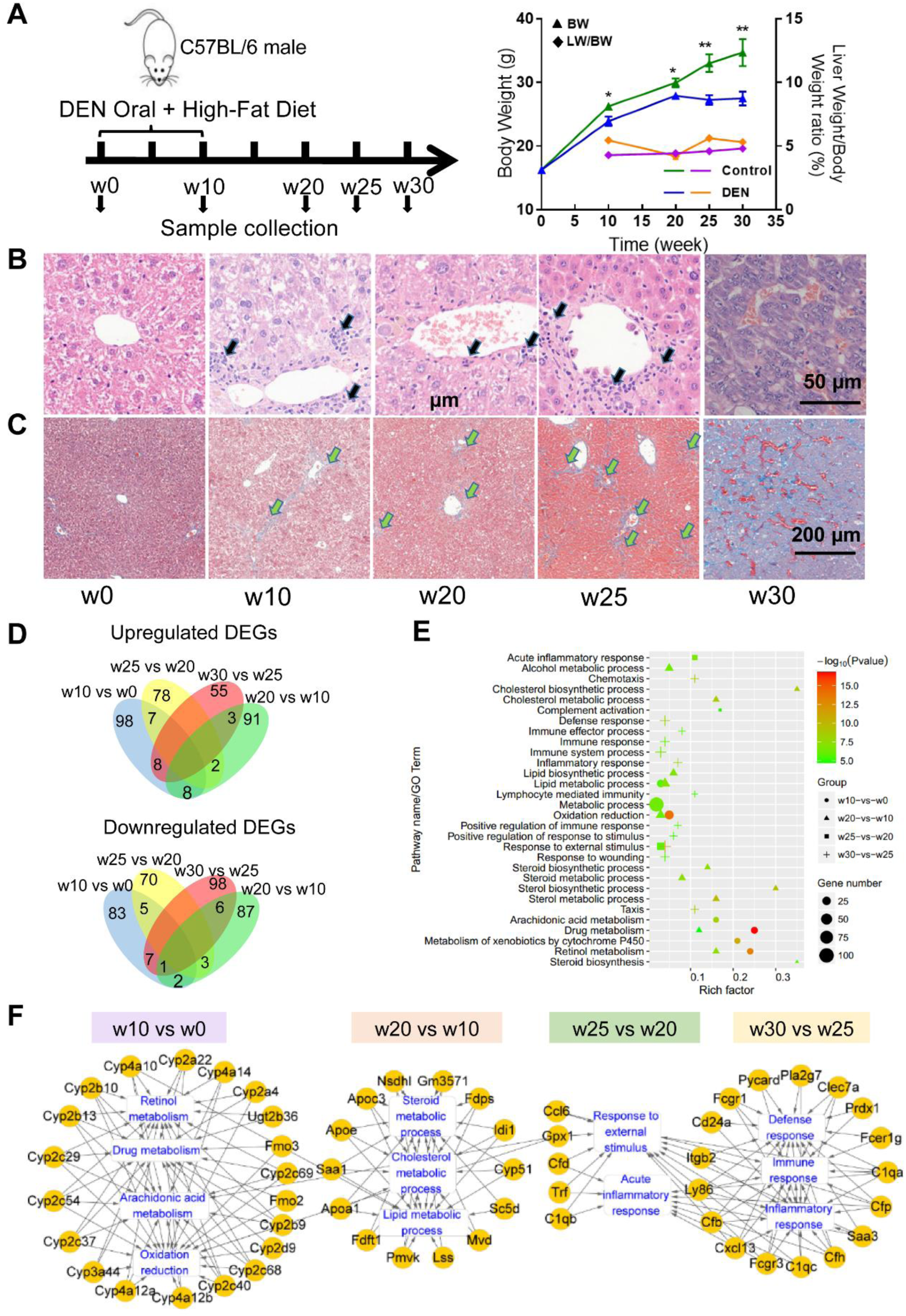
Experimental design, biological characteristics and time series gene expression data for the DEN-induced liver injury model. A Experimental scheme and body weight and liver weight changes. B, C H&E staining and Masson staining of liver tissues at weeks 0, 10, 20, and 25 and tumor tissues at week 30. Black arrow: inflammatory infiltration. Green arrow: fibrosis. D Amount of up- and downregulated DEGs in different comparisons. The upper Venn diagram displays the upregulated DEGs, while the lower Venn diagram displays the downregulated DEGs. E Functional annotation of the DEGs. F Gene-based pathway crosstalk analysis for liver DEGs in the DEN-induced mouse model. *P < 0.05; **P < 0.01.

To identify key regulators in HCC progression, liver samples were collected at five time points with three replicates for each time point, and RNA-seq was performed (Fig 1A). From all samples, we obtained a total of 16,195 expressed genes (fragments per kilobase of transcript per million mapped reads (FPKM)>0). Genes with FPKM values >20 (approximately 10% of the genes) were considered as highly expressed genes. The five groups shared 403 highly expressed genes (Fig EV1A), which were mainly involved in metabolic processes of carboxylic acid, protein and triglyceride, and acute inflammatory response (Fig EV1B).

Gene expression was analyzed to identify key regulators, and 564 DEGs were identified through comparisons between every two adjacent time points (Fig 1D). Functional enrichment analysis revealed that most of the DEGs in the w10 vs w0 comparison were enriched in drug metabolism, retinol metabolism, and oxidation reduction (Fig 1E), indicating metabolic dysfunction in the liver, while DEGs in the w20 vs w10 comparison were mainly enriched in lipid metabolic and biosynthetic pathways, such as cholesterol and steroid. The prominent processes associated with DEGs in the w25 vs w20 comparison were complement activation and inflammatory responses (Fig 1E). Next, we examined the key genes involved in the most significant pathways for each comparison. Through gene-pathway crosstalk analysis, we found that CYP family genes acted as important nodes connecting multiple pathways, including drug metabolism, oxidation reduction, arachidonic acid metabolism and retinol metabolism pathways (Fig 1F). The importance of CYP genes in these pathways has been investigated previously in several studies (Polonikov *et al*, 2017; Ross & Zolfaghari, 2011; Danielson, 2002), and our finding was consistent with the published results.

### Transcriptome analysis and expression validation confirmed *Cyp2c29* as a potential key gene consistently downregulated in HCC development

To further analyze the key gene(s) involved in HCC progression, we screened all 52 CYP genes detected in at least one time point that met a threshold of FPKM > 5.0 (Fig 2A). Approximately half of the CYP genes were continuously downregulated from w0 to w30. This finding is highly relevant to reported findings that P450 genes are downregulated during the inflammatory response (Tsunedomi *et al*, 2005; Xu *et al*, 2001; Wang *et al*, 2018). Most of the downregulated genes belong to the *Cyp2b* and *Cyp2c* subfamilies. Among the identified genes, *Cyp2c29* was highly expressed at w0 but decreased during the liver injury progression (fold change (FC) = −2.88 from w0 to w10, FC = −3.02 from w10 to w20).

**Figure 2.**
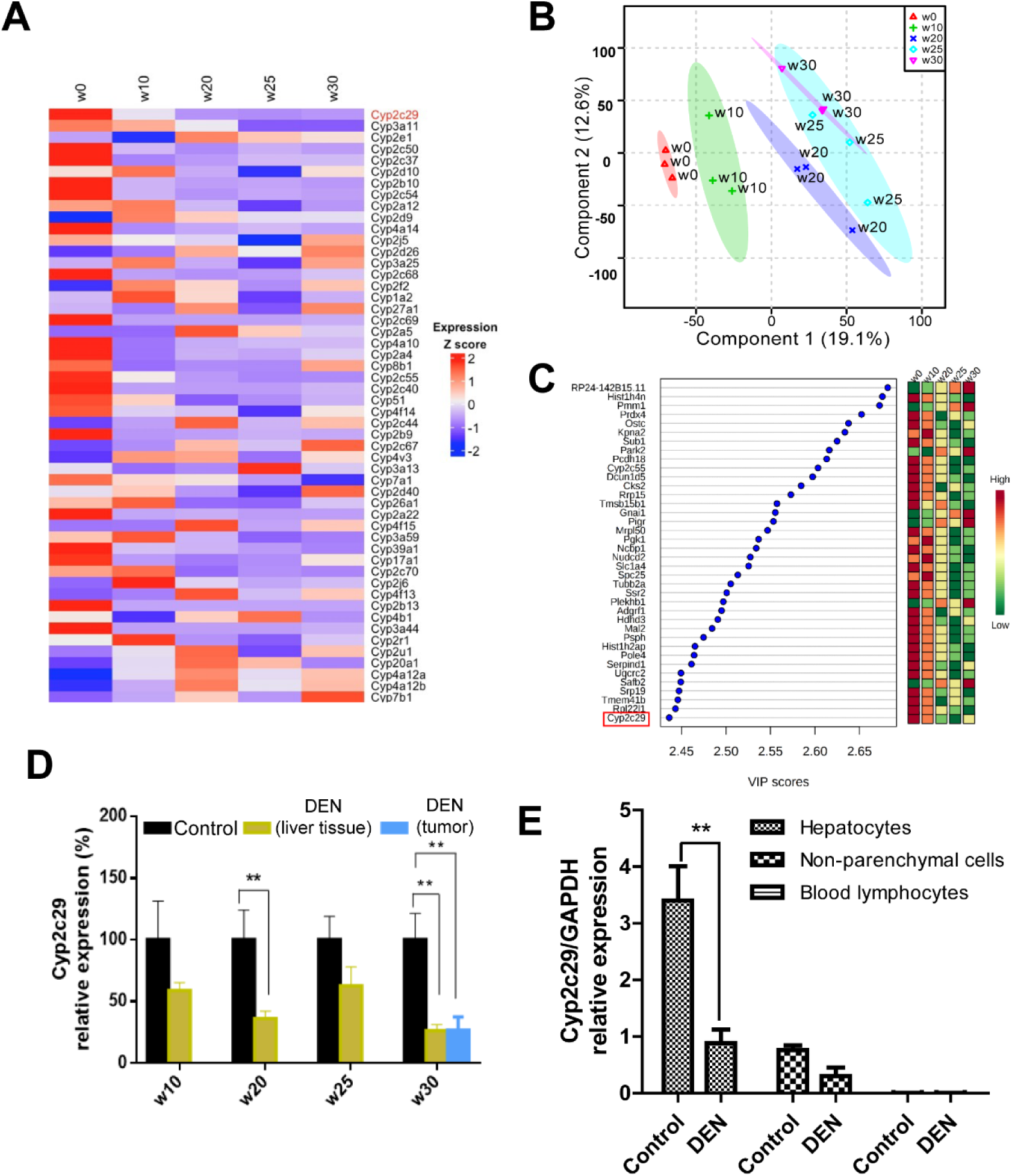
Identification of *Cyp2c29* as a significantly altered gene in DEN-induced liver injury. A Expression changes and clustering of CYP family genes. B, C Important genes identified by PLS-DA and relevant VIP scores. D Validation of decreased *Cyp2c29* expression by qRT-PCR at different time points of DEN-induced liver injury (n=6). E *Cyp2c29* expression in different cells of the injured liver induced by DEN (24 h after 100 mg/kg intraperitoneally injection). **P < 0.01.

In addition, we performed partial least squares discriminant analysis (PLS-DA) of all the genes to identify key genes at each time point. A total of 419 marker genes were obtained with variable importance in projection (VIP) scores >2.0 that contributed to distinguishing among the five groups (Fig 2B). Furthermore, the DEGs with VIP scores in the top 10% were determined (Fig 2C), and *Cyp2c29* was again identified as a core gene. The reduced expression of the *Cyp2c29* gene during liver injury was further confirmed by qRT-PCR analysis (n=6) (Fig 2D). The expression of *Cyp2c29* was lower in the DEN-treated group than that in the control group at the same time point. Hepatocytes constitute about 60-80% of the total cell population in the liver (Racanelli, 2006). We isolated different cells in the liver and found that the expression of the *Cyp2c29* was mainly reduced in the hepatocytes of injured liver induced by DEN (24 h after 100 mg/kg intraperitoneally injection) (Fig 2E).

### *Cyp2c29* suppressed NF-κB activation and inflammation-stimulated cell proliferation

As a member of the Cyp2c family, Cyp2c29 is an endothelial epoxyeicosatrienoic acid (EET) synthase (Sun *et al*, 2010). EETs have anti-inflammatory activity and limit damage to blood vessels caused by inflammation (Tacconelli & Patrignani, 2014). In this study, to investigate the EET’s role in HCC development, we evaluate the effects of EET on hepatocytes under an inflamed environment. To clarify the influence of inflammatory mediators, RAW264.7 cells were treated with lipopolysaccharide (LPS) to mimic an inflamed environment. After LPS treatment, EET reduced the protein levels of p-NF-κB (p65) in RAW264.7 cells (Fig 3A). Moreover, high levels of the inflammatory cytokines in the supernatant (IL-1α, IL-6, IL-17A and MCP-1) secreted by LPS-activated RAW264.7 cells, were reduced after EET treatment (Fig EV3, Fig 3B). The supernatant from LPS-treated RAW264.7 cells significantly enhanced the proliferation viability of primary mouse hepatocytes, whereas decreases of the proliferation viability were shown when combination with EET (Fig 3D), suggesting that Cyp2c29 may suppress cell growth by attenuating inflammation.

**Figure 3.**
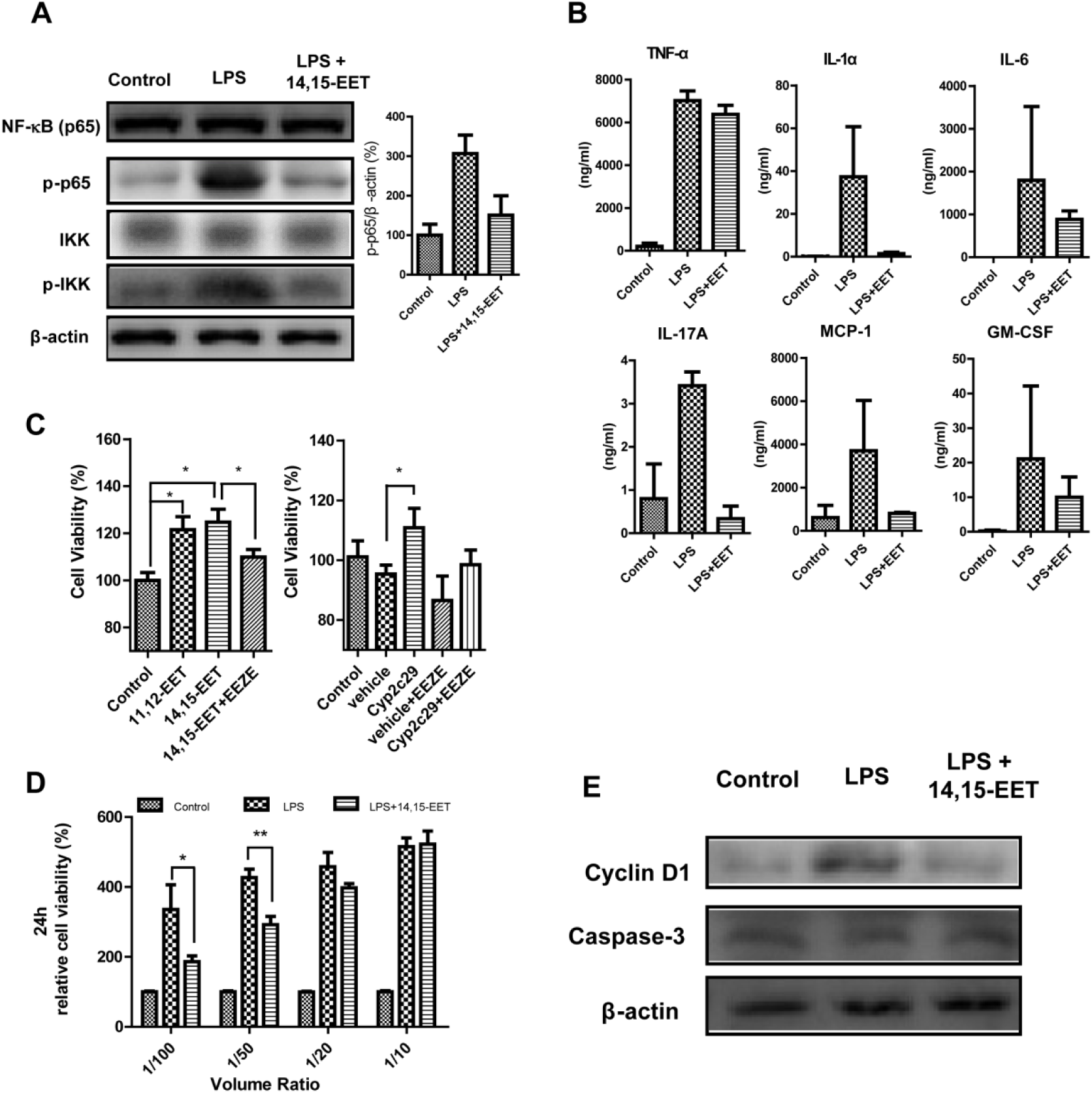
*Cyp2c29* suppressed NF-κB activation and inflammation-stimulated hepatocyte proliferation. A Western blot analysis of NF-κB (p65), p-p65 and p-IKK expression in RAW264.7 ce lls. B Expression of the inflammation related cytokines in the supernatant of macrophage RAW264.7 activated by LPS with or without 14,15-EET treatment. C Cell viability of *Cyp2c29*-transfected or EET-treated HL-7702 cells. D Cell viability of primary mouse hepatocytes incubated with different volume ratios of supernatant from different groups of RAW264.7 cells. E Western blot analysis of cyclin D1 and caspase-3 expression in primary mouse hepatocytes. *P < 0.05; **P < 0.01.

We further investigate the role of Cyp2c29 under normal environment. The liver cell HL-7702 was transfected with a *Cyp2c29* expression plasmid. qRT-PCR results confirmed Cyp2c29 expression (Fig EV2), and EET production analysis further verified it. Significantly enhanced 14,15-EET hydrolysate (14,15-DHET) levels were observed in the supernatant of the Cyp2c29-overexpressing HL-7702 cells (Fig EV2C). HL-7702 cells overexpressing Cyp2c29 or treated with exogenous EETs significantly enhanced cell proliferation viability than the control (Fig 3C), which could be blocked by the EET antagonist 14,15-epoxyeicosa-5(Z)-enoic acid (14,15-EEZE) (Fig 3C). Our result was agreed with the report that EET enhanced cell proliferation *in vitro* (Boron& Boulpaep, 2002). The data suggested that EET plays different roles in cell proliferation with or without inflammatory. Western blot analysis showed that expression of cyclin D1, a critical regulator of cell cycle progression and cell proliferation, was reduced in the EET and LPS co-treatment group, but caspase-3 expression was not significantly changed (Fig 3E). These data indicated that decreased proliferation might be the main cause of the suppressed hepatocyte proliferation viability. Similar results were observed on hepatocellular carcinoma H22 cell line (Fig EV4).

### Overexpression of *Cyp2c29* reduced inflammation in liver injury models

NF-κB is a master regulator of inflammation and is also known to be a central signaling node between hepatic injury and HCC (Luedde & Schwabe, 2011). Consistent with previous reports, we found that the NF-κB signaling pathway was activated during liver inflammation based on data from DEN-induced HCC in mice (Fig EV5). To verify the role of *Cyp2c29* in liver injury, we used two other liver injury models induced by acetaminophen (APAP) or carbon tetrachloride (CCl_4_); these models are characterized by typical NF-κB activation and inflammatory responses (Tan *et al*, 2016; Jiang *et al*, 2017). A *Cyp2c29* expression plasmid was transfected into mice, and the mice were then injected with APAP or CCl_4_ to induce liver injury. The animals were sacrificed at 6 h and 24 h after APAP administration. *Cyp2c29* expression both was increased on mRNA and protein levels in the APAP-treated group transfected with the *Cyp2c29* plasmid (Fig EV6). During pathological examination, we found that the cell necrosis area was reduced in the *Cyp2c29* group, as revealed by H&E staining of liver tissue (Fig 4A-B). Increased alanine aminotransferase (ALT) and aspartate aminotransferase (AST) levels were observed in the APAP group treated with empty vector plasmid, whereas significantly decreased ALT and AST levels were detected in the *Cyp2c29* group (Fig 4C); these results illustrated the protective effect of *Cyp2c29* during liver injury. CCl_4_-treated mice also exhibited significantly lower *Cyp2c29* expression than control mice (Fig EV6), suggested that *Cyp2c29* is consistently downregulated during liver injury. Less inflammatory infiltration and lower ALT levels were observed in the *Cyp2c29* group (transfected with *Cyp2c29* plasmid and treated with CCl_4_) than in the CCl_4_ model group (Fig EV7A-D). In addition, the number of CD68-positive cells in the liver was reduced in the *Cyp2c29* group compared with the CCl_4_ model group (Fig EV7E), which might attribute to decreased inflammatory infiltration and liver injury (McGuinness *et al*, 2000).

**Figure 4.**
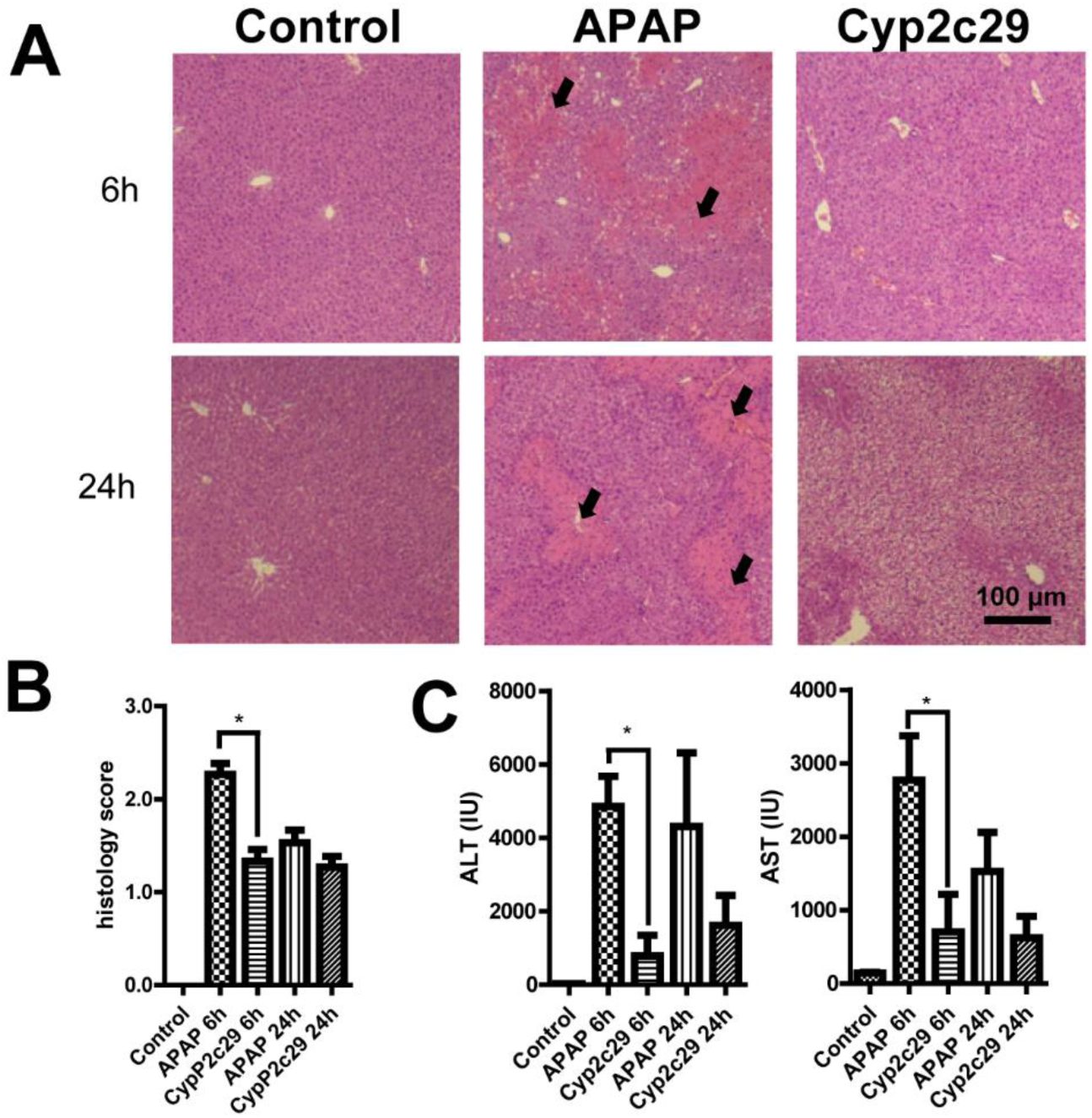
Overexpression of *Cyp2c29* reduced inflammation in the APAP-induced liver injury model. The mice in the *Cyp2c29* group received 100 μg of *Cyp2c29* plasmid DNA 48 h before APAP administration, while those in the APAP group received an equivalent empty vector 48 h before APAP administration. The control group did not receive APAP or plasmid DNA. A, B Liver H&E staining and histology score of APAP-induced liver injury. Black arrows: necrotic areas. C ALT and AST levels at 6 h and 24 h after APAP administration. *P < 0.05.

### Transcriptome analysis and in vivo experiments revealed a potential anti-inflammatory function of *Cyp2c29* in liver injury

To further demonstrate the role of *Cyp2c29* in liver injury, we performed RNA-seq to detect gene expression in APAP models treated with Cyp2c29 plasmid or empty vector plasmid for 6 h or 24 h. We identified 131 upregulated and 261 downregulated DEGs at 6 h, and 81 upregulated and 254 downregulated DEGs at 24 h by comparing the Cyp2c29 treated group with the APAP model group (Fig EV8). Next, the specific functions of the DEGs were investigated (Fig 5A). The results indicated that compared with the control group (at 6 h and 24 h), the APAP model group showed suppressed metabolism and increased inflammation and cell death, while the opposite effects were observed for the APAP group treated with *Cyp2c29*. Cell differentiation and programmed cell death pathways were highly active in the APAP model group but were all suppressed in the Cyp2c29-treated group. These effects were more significant at 24 h than at 6 h after treatment (Fig 5A). We further investigated the TF regulation of the genes in 10 significant pathways. Eleven TFs regulated genes associated with cell proliferation, inflammatory response and metabolism pathways were identified (Fig 5B). Furthermore, we explored the gene expression patterns of these pathways and found that liver tissues in the APAP model group exhibited high inflammatory activity and cell differentiation/death activity at 6 h and 24 h, while such activities were reduced in the Cyp2c29-treated group at 6 h and 24 h (Fig 6A). To further reveal the detailed mechanism, we mapped the DEGs between the Cyp2c29-treated group and APAP model group at 6 h to inflammation-related pathways, including the NF-κB, MAPK, TNF, and IL-17 pathways, and found that these pathways were all suppressed (Fig EV9).

**Figure 5.**
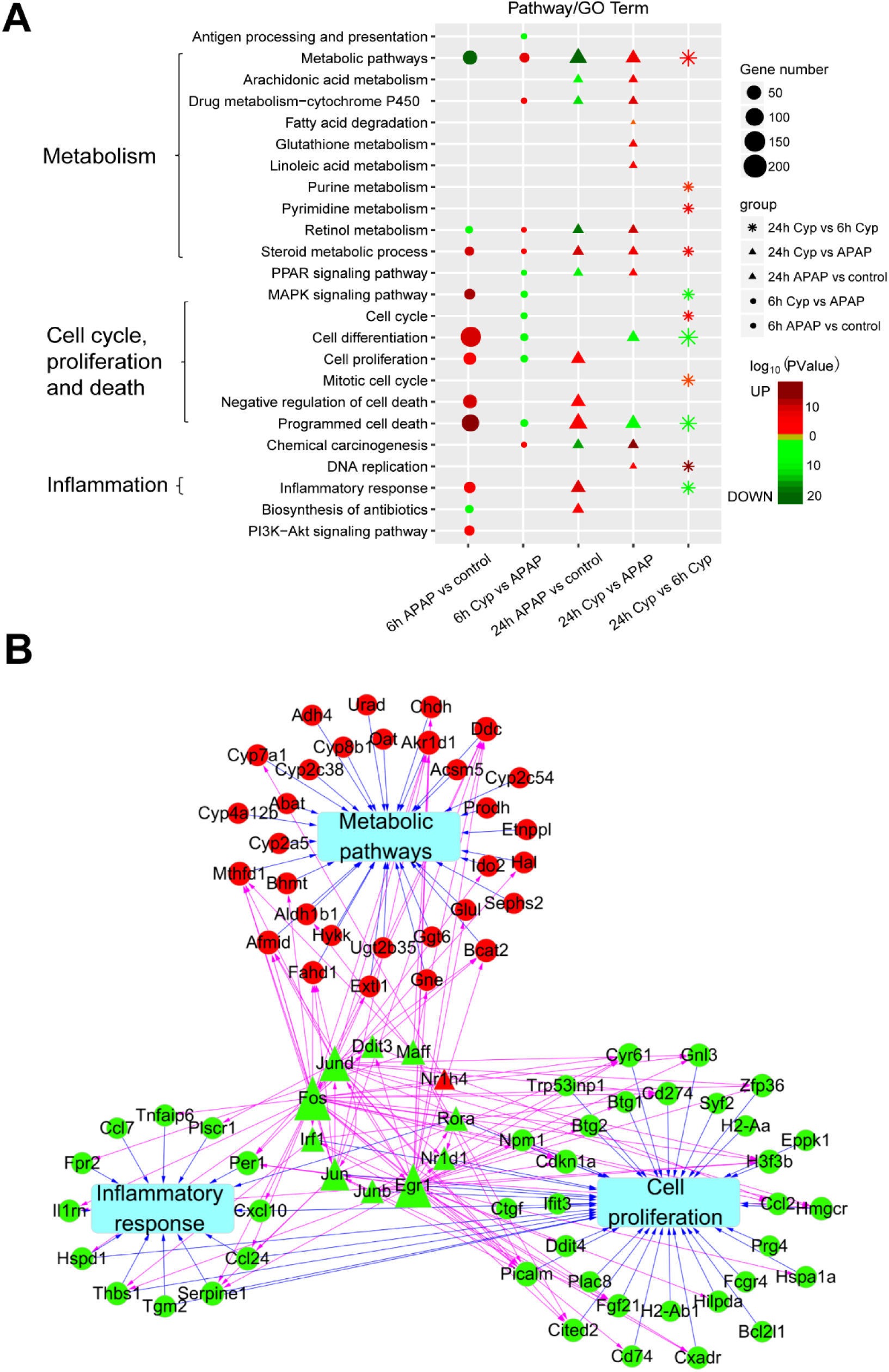
Transcriptome analysis revealed the potential functions of *Cyp2c29* in APAP-induced liver injury. A GO functional enrichment of DEGs from the APAP (model) group vs control group and *Cyp2c29* group vs APAP group comparisons at 6 h and 24 h after APAP treatment. Red: upregulated. Green: downregulated. B TF regulation network and crosstalk for the 6 h *Cyp2c29* group vs APAP group comparison. Red: upregulated. Green: downregulated. Diamonds: TFs. Circles: genes.

**Figure 6.**
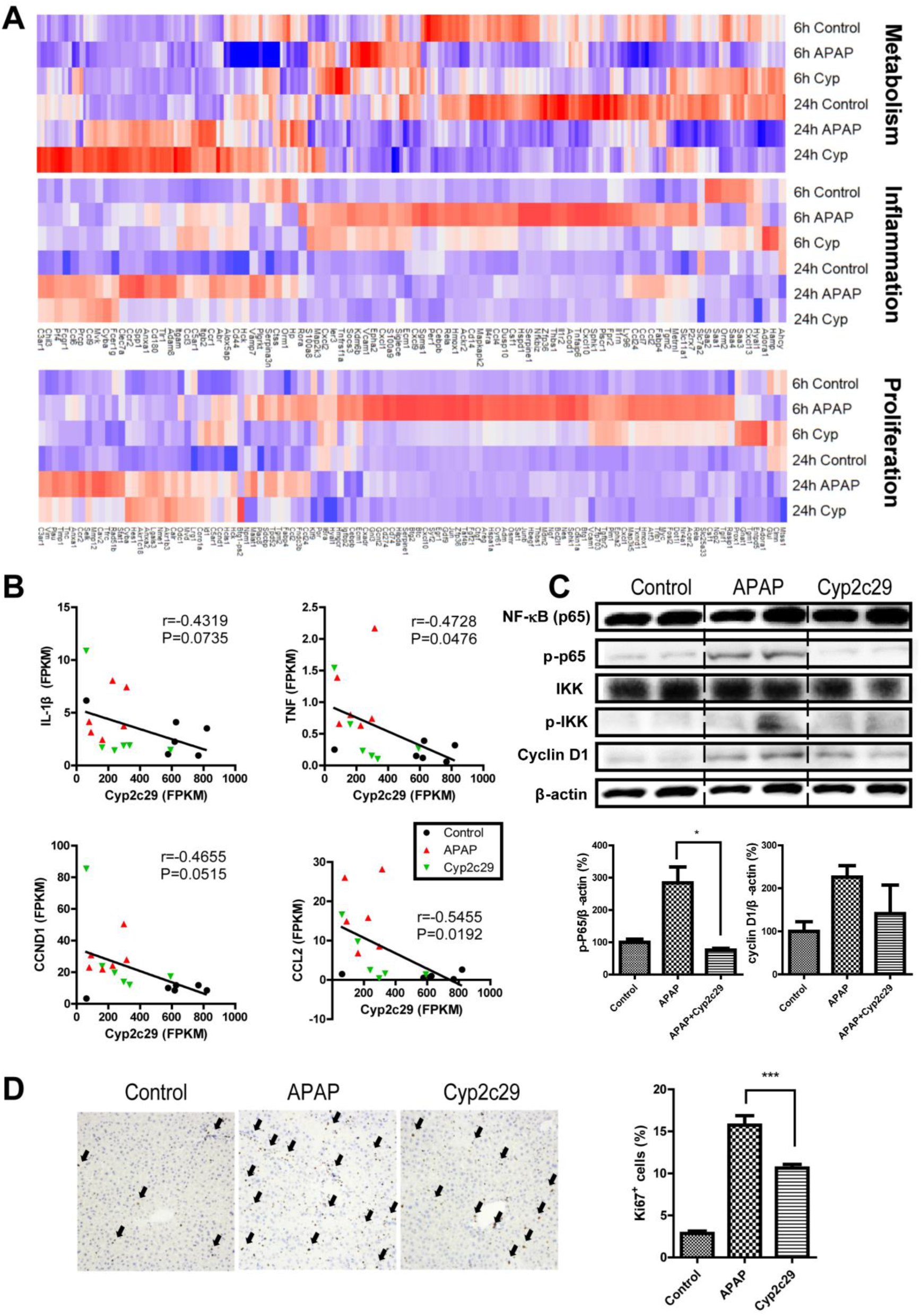
*Cyp2c29* attenuated NF-κB activation and inflammation in APAP-induced liver injury models. A Expression patterns of DEGs associated with metabolism, inflammation and proliferation pathways. B Correlations between *Cyp2c29* expression and IL-1β, TNF, CCND1 and CCL2 expression. C Western blot analysis of p-NF-κB, p-IKK and cyclin D1 expression in APAP-induced liver injury. D Immunohistochemical staining and counts of Ki67-positive cells. Black arrows: Ki67-positive cells. ***P < 0.001.

Notably, we found that the expression of Cyp2c29 was inversely correlated with that of IL-1β, TNF, CCND1 and CCL2 (Fig 6B), which are marker genes on the NF-κB pathway (Taniguchi & Karin, 2018). The protein levels of phosphorylated NF-κB p65 (also known as p-NF-kappaB p65 or p-NF-κB) were lower in the Cyp2c29 group than in the APAP model group (Fig 6C). An upstream regulator of NF-κB, IKK, was phosphorylated and activated in the APAP model group; however, Cyp2c29 overexpression reduced the phospho-IKK levels (Fig 6C). Therefore, Cyp2c29 suppressed the IKK-NF-κB signaling pathway. In the CCl_4_-induced mouse model, decreased p-NF-κB was also observed in the Cyp2c29 group compared with the CCl_4_ model group (Fig EV7F). These results indicated that the NF-κB pathway was suppressed in both the APAP- and CCl_4_-induced liver injury models after pre-treated with the Cyp2c29 plasmid. Immunohistochemical staining of Ki67 showed less liver cell proliferation in APAP-treated Cyp2c29-expressing mice than in APAP model mice (Fig 6D), indicating less compensatory regeneration during liver injury. Consistent with these results, cyclin D1 was confirmed to be downregulated in the Cyp2c29 group (Fig 6C).

### Expression human homologous genes to mouse *Cyp2c29* were positively correlated with survival time in HCC patients

Among human epoxygenases, which are closely associated with lipid metabolism and EET production, the most well-characterized members are CYP2C8, CYP2C9, and CYP2J2 (Bishop-Bailey *et al*, 2014). CYP2C8 and Cyp2c29 share 83.5% protein sequence similarity, while CYP2C9 and Cyp2c29 share 89.0% similarity (Fig 7A). To explore the association between CYP epoxygenase expression and clinical outcomes in HCC patients, we performed an overall survival analysis using gene expression data from 363 HCC patients from the Cancer Genome Atlas (TCGA). The results indicated that high expression of *CYP2C8* and *CYP2C9* was positively correlated with clinical survival (significance: *p* = 0.046 and 0.0097, respectively) (Fig 7B). Next, we quantified the gene expression levels of *CYP2C8* and *CYP2C9* in different HCC stages (TNM stages I, II, III, and IV) according to their clinical information (Fig 7C). The results showed that *CYP2C8* and *CYP2C9* expression significantly declined from stage I to stage III during HCC progression (Kruskal-Wallis, P < 0.001).

**Figure 7.**
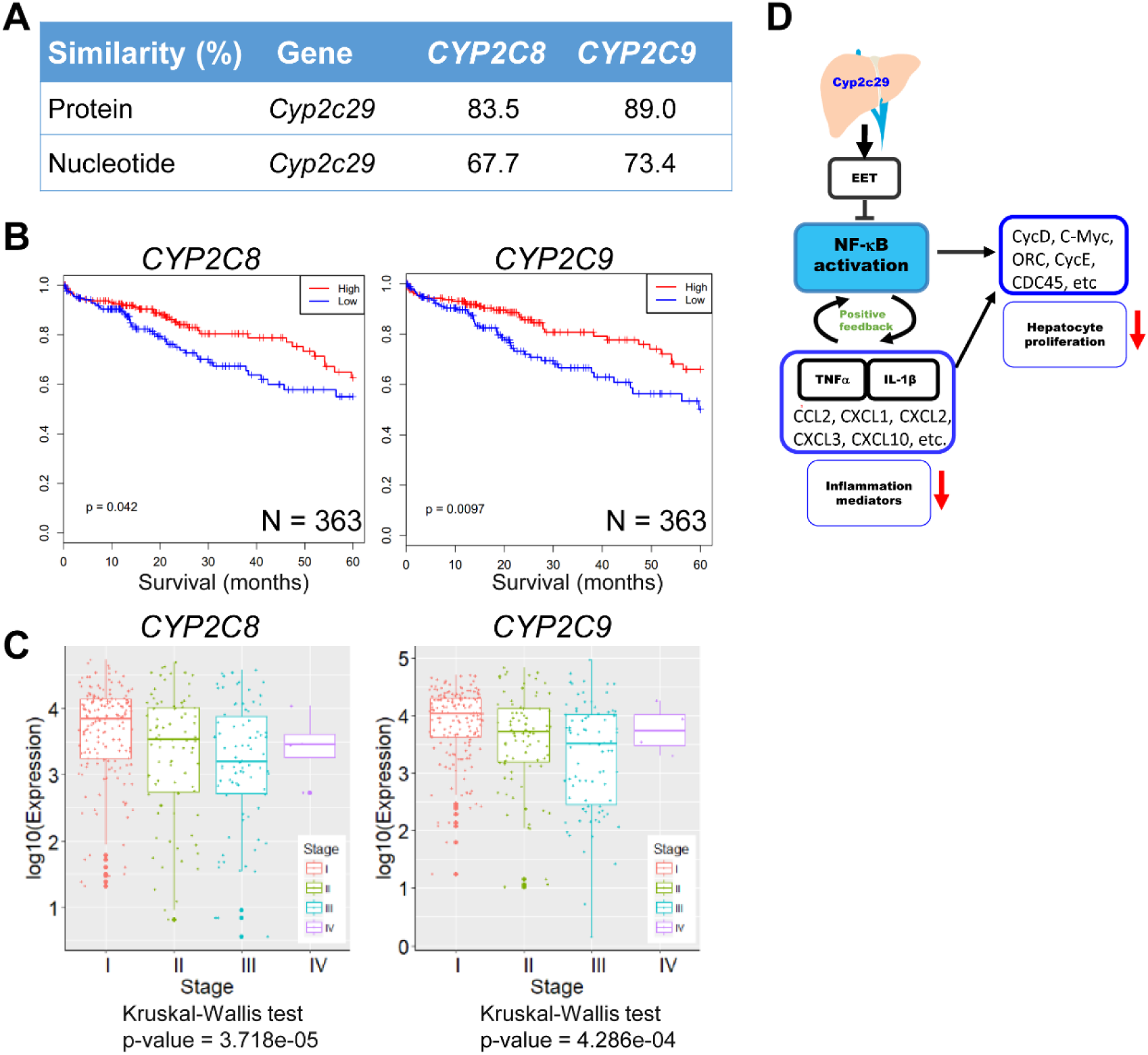
The expression of human homologous genes of mouse *Cyp2c29* is positively correlated with survival time in HCC patients. A Similarities between *Cyp2c29* and human *CYP2C* epoxygenase genes. B Kaplan-Meier plots of 363 HCC patients stratified by *CYP2C8* and *CYP2C9* expression. Expression was classified as high or low based on the median value. C Expression of *CYP2C8* and *CYP2C9* among patients of different cancer stages. D Scheme of the role of *Cyp2c29* in mediating liver inflammation and hepatocyte proliferation through regulation of NF-κB. *Cyp2c29-*derived EET suppresses NF-κB activation and decreases downstream inflammatory pathway gene expression.

## Discussion

As one of the most lethal cancers worldwide, HCC, occurs in the context of chronic liver injury and inflammation (Bishayee, 2014). Pathologic changes, including inflammatory infiltration, regeneration, fibrosis, and tumorous nodules, were observed in our DEN-induced mouse model, which are similar to changes associated with human HCC progression. Based on this model and time series gene expression analysis, we revealed *Cyp2c29* as a key gene in the progression from liver injury to HCC and further characterized its function.

*Cyp2c29* has been confirmed to be a specific CYP isoform responsible for EET-mediated vasodilator responses in mice (Sun *et al*, 2010); however, its role in HCC progression remains largely unknown. In our study, we found that the proliferation of primary hepatocytes was enhanced by supernatant from LPS-activated RAW264.7 cells, whereas the results were reversed by 14,15-EET, consistent with a previous report of the anti-inflammatory activity of 14,15-EET (Theken *et al*, 2011; Schmelzer *et al*, 2005; Oni-Orisan *et al*, 2013). It has been reported that increases in EETs reduce vascular smooth muscle cell proliferation through downregulation of cyclin D1 (Davis *et al*, 2002). Consistent with this report, our results showed that cyclin D1 expression was decreased while 14,15-EET was enhanced under conditions mimicking an inflammatory environment. Collectively, the production of EETs could be part of the reason why *Cyp2c29* suppresses compensatory proliferation during liver injury.

The inflammatory reaction is commonly correlated with HCC progression. (Karin & Lin, 2002; Iimuro *et al*, 1998). Our results showed that the NF-κB pathway was activated in the DEN-treated mouse model. APAP- and CCl_4_-induced liver injury induced an acute inflammatory response that was not similar to the DEN-induced inflammation in our study. However, APAP- and CCl_4_-induced liver injury models are characterized by typical inflammatory responses and NF-κB pathway activation (Ruepp *et al*, 2002; Son *et al*, 2007). Therefore, APAP- and CCl_4_-induced liver injury models were used to verify the role of *Cyp2c29.* In this study, mice transfected with the *Cyp2c29* plasmid showed reduced injury and inflammation in both the APAP and CCl_4_ models, suggesting a protective effect of *Cyp2c29* against liver injury. When *Cyp2c29* was overexpressed, the levels of p-NF-κB and the expression of genes on NF-κB-related pathways, such as TNF and MAPK, which play important roles in inflammation and cell proliferation (Wagner & Nebreda, 2009; Grivennikov & Karin, 2011), were downregulated. NF-κB is critical in the regulation of liver disease, influencing hepatocyte survival, Kupffer cell and hepatic stellate cell activation (Luedde & Schwabe, 2011). NF­κB activation is induced by infection, necrotic cell products, oxidative stress, and inflammatory cytokines such as TNF and IL-1 (Taniguchi & Karin, 2018). Moreover, NF-κB activation leads to mediate the synthesis of proinflammatory cytokines (such as TNF, IL•1 and IL•6) and chemokines (such as CXCL1, CXCL2, and CCL2) (Tak & Firestein, 2001). NF-κB and its related inflammatory molecules form a positive feedback loop and stimulate hepatocyte proliferation and liver regeneration, which may increase the risk of HCC. In this study, we found that *Cyp2c29* overexpression significantly downregulate NF-κB signaling. We performed network analysis and illustrated the involvement of *Cyp2c29* in signaling pathways (Fig 7D). Genes that encode inflammatory cytokines (such as TNF and IL•1), chemokines (such as CCL2, CXCL1, CXCL2, and CXCL3) and cell proliferation-related proteins (such as CycD and CycE) were downregulated when *Cyp2c29* was overexpressed in the mouse model of liver injury. These results demonstrate that Cyp2c29 may reduce liver injury through suppression of the inflammatory response.

High expression levels of *CYP2C8* and *CYP2C9*, highly homologous to *Cyc2C29*, were found to be positively correlated with survival time in HCC patients. Reduced expression levels of *CYP2C8* and *CYP2C9* were observed from stage I to stage III in HCC patients. However, another homologous gene *CYP2J2* expression was not significantly associated with survival time in HCC patients from TCGA (Fig EV10), although CYP2J2 expression has been reported to be associated with tumor progression (Jiang *et al*, 2005; Panigrahy *et al*, 2010). This finding suggests that the role of CYP2J2 may vary in different cancers. These data together indicate the important roles of Cyp2c29 in liver injury and HCC progression.

In conclusion, Cyp2c29 was downregulated during liver injury in different mouse models. Cyp2c29 overexpression enhanced 14,15-EET production and suppressed inflammation-induced hepatocyte proliferation by inhibiting the IKK-NF-κB pathway during liver injury. Our data provide evidence that CYP2C epoxygenases could be potential therapeutic targets for liver diseases. Future studies investigating the effects of other epoxygenases in the CYP2C family may elucidate additional regulators in HCC progression.

## Materials and Protocols

### Mice

Four-week-old male C57BL/6 mice were purchased from the Hubei Research Center of Laboratory Animals (Wuhan, China; quality certification number: SCXK (E) 2015-0018). The animal studies were approved by the Ethics Committee on Animal Experimentation of Huazhong University of Science and Technology (Wuhan, China) and abided by the guidelines of the Science and Technology Department of Hubei Province. All mice were treated with humane care. They were maintained in individual ventilated cages, given autoclaved water and fed irradiated food.

### DEN-induced liver injury model

To mimic the process of HCC development, mice were orally administered diethylnitrosamine (DEN) (TCI Japan) at a dose of 0.16 mmol·kg^−1^ body weight once a week for the first ten weeks and fed a high fat diet (HFD) for 30 weeks (Park *et al*, 2010; Bakiri &Wagner). In the control group, mice were administered sesame oil (the solvent of DEN) once a week for the first ten weeks and fed a basal chow diet. Separately at weeks 0, 10, 20, 25 and 30, mice were sacrificed, and liver tissues were collected for histological examination and RNA extraction.

### Histological examination

Liver tissues were fixed in 4% formaldehyde for 24 h. Paraffin sections (5 μm) were prepared for H&E staining for general morphological observation. Masson trichrome staining was used to reveal collagen fibers. The sections were observed using a light microscope (Nikon Eclipse TE2000-U) and then photographed at 100×, 200×, or 400× magnification. Liver histology scoring was performed as described previously (Hoque *et al*, 2012).

### Sequencing and identification of differentially expressed genes

High-throughput RNA-seq techniques were used to detect gene expression. Three biological replicates for each group were sequenced. For RNA-seq, ribosomal RNA was first removed, and 150 bp paired-end sequencing was carried out on an Illumina HiSeq 3000 platform. All of the RNA isolation, sequencing and data filtering procedures were performed by RiboBio (Guangzhou, China). Assembly, alignment, quantification and profiling of the RNA-seq data were performed on our own server. First, the quality of the reads was verified with FastQC (version: 0.11.3). Then, HISAT2 (version: 2.0.5) was used to map the sequencing reads to the mouse genome (GRCm38). Subsequently, StringTie (version: 1.2.2) was used to assemble the alignments into potential transcripts according to the reference sequences (Pertea *et al*, 2016). Finally, transcript abundance was calculated as the FPKM. Additionally, NOISeq was used to identify DEGs with thresholds of a false discovery rate (FDR) < 0.01 and an |FC| ≥ 1.5. The data were submitted to BIG Data Center (http://bigd.big.ac.cn), and the assigned accession number of the submission is CRA000931.

### Selection of key genes and pathways

For functional annotation, DEGs enrichment analysis was performed with the Database for Annotation, Visualization and Integrated Discovery (DAVID; https://david.ncifcrf.gov/tools.jsp). The Gene Ontology (GO) terms and Kyoto Encyclopedia of Genes and Genomes (KEGG) pathways are displayed in bubble plots. The rich factor in the bubble plots was calculated as the ratio of the number of genes that mapped to a certain pathway to the total number of genes in that pathway. Gene expression patterns were identified by hierarchical clustering and are displayed using a heat map. Furthermore, we investigated the most important pathways (Benjamini-Hochberg adjusted P < 0.05) and their crosstalk. For key gene selection, genes in the top-ranked pathways (P < 0.01) were deemed important. Then, we focused on the genes that were expressed in more than 80% of the groups. PLS-DA was employed to screen the important genes that contributed the most to differentiation of the groups. The cutoff was set as a VIP score >2.0 (the VIP score is a weighted sum of squares of the PLS loadings) (Xia *et al*, 2015).

### *Cyp2c29* plasmid transfection

The *Cyp2c29* pENTER-C-GFP plasmid and plasmid-harboring *E. coli* were supplied by Vigene Bioscience. The plasmid was sequenced to confirm its accuracy (GenBank accession number NM_007815.3). Then, plasmids were extracted and purified from *E. coli* with an EndoFree Plasmid Kit (Tiangen, China). Plasmids were transfected into HL-7702 cells according to the protocol of Lipofectamine^®^ 3000 Transfection Reagent (Thermo Fisher, USA). Hydrodynamics-based transfection of *Cyp2c29* plasmids into mouse liver was performed as described previously (Liu *et al*, 1999).

### Treatment of HL-7702 cells with EET

The hepatocyte line HL-7702 was transfected with *Cyp2c29* plasmid or treated with exogenous 11,12-EET or 14,15-EET (Cayman, USA) at a concentration of 100 nmol/L (Shen *et al*, 2008). Cell viability was assessed with a Cell Counting Kit-8 (Dojindo, Japan). The levels of a stable metabolite of 14,15-EET, 14,15-DHET, in the cell supernatant were measured with a 14,15-DHET ELISA kit (Detroit R&D, USA) according to the manual.

### Isolation of hepatocytes and non-parenchymal cells

Purification of primary hepatocytes was performed as described previously (Amaral *et al*, 2013). Hepatocytes were separated with 90% Percoll solution (Aparicio-Vergara *et al*, 2017), and non-parenchymal cells were separated with 20 % iodixanol solution, separately (Aparicio-Vergara *et al*, 2015).

### Incubation of primary mouse hepatocytes with LPS-activated RAW264.7 cell supernatant

RAW264.7 cells were activated with 200 ng/mL LPS (Sigma-Aldrich, USA) with or without 100 nmol/L 14,15-EET added to the medium (Shen *et al*, 2008). The supernatant was collected 12 h later. Cytokines in the supernatant were quantified by LEGENDplex™ Mouse Inflammaton Panel (BioLegend, USA). A series of RAW264.7 supernatants with volume ratios ranging from 1/100 to 1/10 were mixed into the primary mouse hepatocyte medium. Cell viability was assessed with a Cell Counting Kit-8 (Dojindo, Japan) after 24 h of incubation.

### APAP- and CCl_4_-induced liver injury models

APAP- and CCl_4_-induced mouse liver injury models were established by intraperitoneal injection of mice with 500 mg/kg APAP and 10 ml/kg CCl_4_ (0.2%, dissolved in olive oil), respectively. The mice were divided into 3 groups in each experiment: a control group (n = 6), a model group (100 μg of empty plasmid was diluted to 0.1 ml/g body weight), and a *Cyp2c29* group (100 μg of *Cyp2c29* plasmid was diluted to 0.1 ml/g body weight). Forty-eight hours post plasmid injection, APAP and CCl_4_ were administered to the *Cyp2c29* groups and model groups. Mice were sacrificed at 6 h and 24 h after APAP administration (or 24 h and 48 h after CCl_4_ administration) for collection of serum and liver tissues. Serum ALT and AST levels were determined with an automatic biochemical analyzer (Beckman, Germany) to determine the extent of hepatocyte injury. Hepatocyte proliferation was measured by Ki67 staining. The data are expressed as the mean ± standard error of the mean (SEM). RNA extraction and high-throughput sequencing were performed by BGI (Wuhan, China). Gene expression, DEGs and gene enrichment were analyzed by the same methods as those described above for the time series gene expression analysis. For the TF regulatory network analysis, we used the same method described in our previous publication (Zhang *et al*, 2015). Finally, Cytoscape (version 3.2.1) was used to visualize the networks. The data were submitted to BIG Data Center(http://bigd.big.ac.cn), and the assigned accession number of the submission is CRA000931.

### Statistical analysis

In the APAP experiment and the CCl_4_ experiment, Bonferroni and Dunnett T3 tests were used for multiple comparisons after ANOVA based on homoscedasticity, respectively. A significance level of 0.05 and a power of 0.9 were used to estimate the necessary sample size. Based on our experimental design and pre-experiments, the difference between the means divided by the pooled standard deviation (effect size) was greater than 3; n=3.370694 was obtained for each group with the pwr.t.test function in the R programming language. Ultimately, a sample size of 6 for each group was selected to ensure the robustness of our experiments. Statistical analysis was performed using the software program IBM SPSS Statistics (version 20). Statistical significance was determined by Student’s t test for the other experiments.

### qRT-PCR

Total RNA from hepatic tissues or cells was extracted using TRIzol reagent (Thermo Fisher, USA). RNA was reverse-transcribed into cDNA with a reverse transcription kit (Takara, Japan), and qRT-PCR was performed with a SYBR^®^ Premix Ex Taq II Kit (Takara, Japan) and a 7500 Real-Time PCR System (Applied Biosystems). The GAPDH gene was selected as the housekeeping gene.

### Western blot analysis

To enable loading of equal amounts, protein concentrations were determined with a bicinchoninic acid (BCA) protein assay reagent kit (Beyotime, China). The protein samples were adjusted to the same concentration before loading. HL-7702 cell and liver tissue protein extracts were fractionated by electrophoresis on 12% SDS-polyacrylamide gels (Beyotime, China) and then transferred to nitrocellulose membranes. The protein blots were blocked with 5% BSA dissolved in Tris-buffered saline with Tween 20 and then incubated with appropriate primary antibodies against NF-κB p65 (1:500; Proteintech, 10745-1-AP), p-NF-κB p65 (1:500; Cell Signaling Technology, #3031), cyclin D1 (1:500; Cell Signaling Technology, #2978), IKK (1:1000; Proteintech, 15649-1-AP), phospho-IKK (1:500; Cell Signaling Technology, #2694), cleaved caspase-3 (1:1000; Cell Signaling Technology, #9664), GAPDH (1:1000; Proteintech, 60004-1-1g) and β-actin (1:2000; Cell Signaling Technology #8457) at 4°C for 12 h. After that, the membranes were washed three times and incubated with a 1:3000 dilution of horseradish peroxidase-conjugated secondary antibody (Yeasen, China) for 2 h. Then, the proteins were detected with an enhanced chemiluminescence (ECL) system (Bio-Rad, USA).

### Targeted proteomics

A quantitation of liver Cyp2c29 protein in mice after APAP and CCl_4_ treatment was performed by targeted proteomics according to the previous reports (Kockmann *et al*, 2016). Tissue samples were lysed in a solution containing 200 μl of thiourea buffer (50 mM Tris-Cl, pH 8, 7 M urea, 2 M thiourea, and 1 × protease inhibitors cocktails), 800 μl of ice-cold acetone, and 10 mM dithiothreitol. After centrifugation at 13,000×g for 20 min at 4 °C, the precipitated pellets were washed 3 times and collected. The dried pellets were dissolved in 200 μl of thiourea buffer for further analysis. Triple TOF 5600+ LC-MS/MS system (AB SCIEX) was applied for Targeted MS analysis using parallel reaction monitoring (PRM). Spectra library generation was analyzed by ProteinPilot software. The selected peptides for quantification were according to the ion signals in spectra library.

### Survival analysis in human HCC

To verify the impact of *Cyp2c29* homologous genes in humans, we investigated the effects of homologous gene expression on clinical survival. The homologous genes were obtained from HomoloGene (https://www.ncbi.nlm.nih.gov/homologene/), and their similarities and identities were calculated with Needle from EMBOSS (version: 6.6.0). The gene expression data of 363 patients with clinical survival information were downloaded from TCGA (Uhlén *et al*, 2015). Kaplan-Meier analysis and log-rank tests were used for survival analysis as in GSCALite (http://bioinfo.life.hust.edu.cn/web/GSCALite/) (Liu *et al*, 2018). The samples were divided into high expression groups and low expression groups for each gene based on the median expression of the 363 patients. A value of *P* < 0.05 was considered significant. Subsequently, the patients were divided into four groups according to their clinical information (by TNM stage: I, II, III, and IV), and the CYP gene expression at different stages is shown with a boxplot. The normality of the distribution was examined, and significance was tested with the Kruskal-Wallis test.

## Abbreviations

HCC: hepatocellular carcinoma
DEN: diethylnitrosamine
APAP: acetaminophen
EET: epoxyeicosatrienoic acid
ALT: alanine aminotransferase
AST: aspartate transaminase
DEG: differentially expressed gene
CYP: cytochrome P450
LPS: lipopolysaccharide

## Acknowledgements

This study is supported by The National Key Research and Development Program of China (2017YFA0700403), National Natural Science Foundation of China (Nos.81573013, 31822030, and 31771458), National Basic Research Program of China (2018YFA0208903).

## Author Contributions

An-Yuan Guo, Yanhong Zhu and Xiangliang Yang designed the study. Qi Wang, Lijun Zhao and Yuxin Wu performed animal experiments. Qin Tang, Qiong Zhang, Hui Hu and Lan-Lan Liu performed the bioinformatics analysis. Qi Wang, Qin Tang, Xiang Liu, Yanhong Zhu and An-Yuan Guo wrote the manuscript.

## References

Amaral SS, Oliveira AG, Marques PE, Quintão JL, Pires DA, Resende RR, Sousa BR, Melgaço JG, Pinto MA, Russo RC et al (2013) Altered responsiveness to extracellular ATP enhances acetaminophen hepatotoxicity. Cell Commun Signal 11: 10.

Aparicio-Vergara M, Tencerova M, Morgantini C, Barreby E, Aouadi M (2017) Isolation of Kupffer cells and hepatocytes from a single mouse liver. Methods Mol Biol 1639: 161–171.

Aparicio-Vergara M, Tencerova M, Morgantini C, Barreby E, Aouadi M (2015) Isolation of non-parenchymal cells from the mouse liver. Methods Mol Biol 1325: 3–17.

Bakiri L, Wagner EF (2013) Mouse models for liver cancer. Mol Oncol 7: 206–223.

Balkwill F, Mantovani A (2001) Inflammation and cancer: back to Virchow? Lancet 357: 539–545.

Bishayee A (2014) The role of inflammation and liver cancer. Advances in experimental medicine and biology (Inflammation and Cancer) 816: 401–435.

Bishop-Bailey D, Thomson S, Askari A, Faulkner A, Wheeler-Jones C (2014) Lipid-metabolizing CYPs in the regulation and dysregulation of metabolism. Annu Rev Nutr 34: 261–279.

Boron WF, Boulpaep EL (2002) Medical physiology: a cellular and molecular approach. Saunders, an imprint of Elsevier Science.

Colombo M, Maisonneuve P. Controlling liver cancer mortality on a global scale: Still a long way to go (2017) J Hepatol 67: 216–217.

Danielson PB (2002) The cytochrome P450 superfamily: Biochemistry, evolution and drug metabolism in humans. Curr Drug Metab 3: 561–597.

Davis BB, Thompson DA, Howard LL, Morisseau C, Hammock BD, Weiss RH (2002) Inhibitors of soluble epoxide hydrolase attenuate vascular smooth muscle cell proliferation. Proc Natl Acad Sci USA 99: 2222–2227.

Ferlay J, Soerjomataram I, Dikshit R, Eser S, Mathers C, Rebelo M, Parkin DM, Forman D, Bray F (2015) Cancer incidence and mortality worldwide: Sources, methods and major patterns in GLOBOCAN2012. Int J Cancer 136: E359–E86.

Grivennikov SI, Karin M (2011) Inflammatory cytokines in cancer: tumour necrosis factor and interleukin 6 take the stage. Ann Rheum Dis 70: i104–108.

He G, Karin M (2011) NF-kappa B and STAT3 - key players in liver inflammation and cancer. Cell Res 21:159–168.

Hoque R, Sohail MA, Salhanick S, Malik AF, Ghani A, Robson SC, Mehal WZ (2012) P2X7 receptor-mediated purinergic signaling promotes liver injury in acetaminophen hepatotoxicity in mice. Am J Physiol Gastrointest Liver Physiol 302: G1171–1179.

Iimuro Y, Nishiura T, Hellerbrand C, Behrns KE, Schoonhoven R, Grisham JW, Brenner DA (1998) NF kappa B prevents apoptosis and liver dysfunction during liver regeneration. J Clin Invest 101: 802–811.

Jiang JG, Chen CL, Card JW, Yang S, Chen JX, Fu XN, Ning YG, Xiao X, Zeldin DC, Wang DW (2005) Cytochrome P450 2J2 promotes the neoplastic phenotype of carcinoma cells and is up-regulated in human tumors. Cancer Res 65: 4707–4715.

Jiang L, Yue S, Li C, Ke M, Busuttil RW, Kupiec-Weglinski JW, Li J, Ying QL, Ke B (2017) TAK1 signaling is essential for NLRP3 activation in mouse drug-induced damage-associated hepatitis. J Hepatol 66: S396.

Karin M, Lin A (2002) NF-kappa B at the crossroads of life and death. Nat Immunol 3: 221–227.

Kockmann T, Trachsel C, Panse C, Wahlander A, Selevsek N, Grossmann J, Wolski WE, Schlapbach R (2016) Targeted proteomics coming of age - SRM, PRM and DIA performance evaluated from a core facility perspective. Proteomics 16: 2183–2192.

Li H, Clarke JD, Dzierlenga AL, Bear J, Goedken MJ, Cherrington NJ (2017) In vivo cytochrome P450 activity alterations in diabetic nonalcoholic steatohepatitis mice. J Biochem Mol Toxic. 31.

Lin Y, Sibanda VL, Zhang HM, Hu H, Liu H, Guo AY (2015) MiRNA and TF co-regulatory network analysis for the pathology and recurrence of myocardial infarction. Sci Rep 5: 9653.

Lin Y, Zhang Q, Zhang HM, Liu W, Liu CJ, Li Q, Guo AY (2015) Transcription factor and miRNA co-regulatory network reveals shared and specific regulators in the development of B cell and T cell. Sci Rep 5: 15215.

Liu CJ, Hu FF, Xia MX, Han L, Zhang Q, Guo AY (2018) GSCALite: a web server for gene set cancer analysis. Bioinformatics 34: 3771–3772.

Liu F, Song Y, Liu D (1999) Hydrodynamics-based transfection in animals by systemic administration of plasmid DNA. Gene Ther 6: 1258–1266.

Luedde T, Schwabe RF (2011) NF-κB in the liver-linking injury, fibrosis and hepatocellular carcinoma. Nat Rev Gastro Hepat 8: 108–118.

McGuinness PH, Painter D, Davies S, McCaughan GW (2000) Increases in intrahepatic CD68 positive cells, MAC387 positive cells, and proinflammatory cytokines (particularly interleukin 18) in chronic hepatitis C infection. Gut 46: 260–269.

Nebert DW, Dalton TP (2006) The role of cytochrome P450 enzymes in endogenous signalling pathways and environmental carcinogenesis. Nat Rev Cancer 6: 947–960.

Oni-Orisan A, Deng Y, Schuck RN, Theken KN, Edin ML, Lih FB, Molnar K, DeGraff L, Tomer KB, Zeldin DC et al (2013) Dual modulation of cyclooxygenase and CYP epoxygenase metabolism and acute vascular inflammation in mice. Prostaglandins Other Lipid Mediat 104: 67–73.

Panigrahy D, Kaipainen A, Greene ER, Huang S (2010) Cytochrome P450-derived eicosanoids: the neglected pathway in cancer. Cancer Metastasis Rev 29: 723–735.

Park EJ, Lee JH, Yu GY, He G, Ali SR, Holzer RG, Osterreicher CH, Takahashi H, Karin M (2010) Dietary and genetic obesity promote liver inflammation and tumorigenesis by enhancing IL-6 and TNF expression. Cell 140: 197–208.

Pertea M, Kim D, Pertea GM, Leek JT, Salzberg SL (2016) Transcript-level expression analysis of RNA-seq experiments with HISAT, StringTie and Ballgown. Nat Protoc 11: 1650–1667.

Polonikov A, Bykanova M, Ponomarenko I, Sirotina S, Bocharova A, Vagaytseva K, Stepanov V, Churnosov M, Bushueva O, Solodilova M et al (2017) The contribution of CYP2C gene subfamily involved in epoxygenase pathway of arachidonic acids metabolism to hypertension susceptibility in Russian population. Clin Exp Hypertens 39: 306–311.

Racanelli V, Rehermann B (2006) The liver as an immunological organ. Hepatology 43: S54–S62.

Ray K (2018) A complex interplay between inflammation and immunity in liver cancer. Nat Rev Gastro Hepat 15: 3.

Ross AC, Zolfaghari R (2011) Cytochrome P450s in the regulation of cellular retinoic acid metabolism. Annu Rev Nutr 31: 65–87.

Ruepp SU, Tonge RP, Shaw J, Wallis N, Pognan F (2002) Genomics and proteomics analysis of acetaminophen toxicity in mouse liver. Toxicol Sci 65: 135–150.

Schmelzer KR, Kubala L, Newman JW, Kim IH, Eiserich JP, Hammock BD (2005) Soluble epoxide hydrolase is a therapeutic target for acute inflammation. Proc Natl Acad Sci USA 102: 9772–9777.

Schuck RN, Zha W, Edin ML, Gruzdev A, Vendrov KC, Miller TM, Xu Z, Lih FB, DeGraff LM, Tomer KB et al (2014) The Cytochrome P450 epoxygenase pathway regulates the hepatic inflammatory response in fatty liver disease. Plos One 9: e110162.

Shen GF, Jiang JG, Fu XN, Wang DW (2008) Promotive effects of epoxyeicosatrienoic acids (EETs) on proliferation of tumor cells [Article in Chinese]. Ai Zheng 27: 1130–1136.

Son G, Iimuro Y, Seki E, Hirano T, Kaneda Y, Fujimoto J (2007) Selective inactivation of NF-kappa B in the liver using NF-kappa B decoy suppresses CCl_4_-induced liver injury and fibrosis. Am J Physiol Gastrointest Liver Physiol 293: G631–G639.

Sun D, Yang YM, Jiang H, Wu H, Ojaimi C, Kaley G, Huang A (2010) Roles of CYP2C29 and RXR gamma in vascular EET synthesis of female mice. Am J Physiol Regu. Integr Comp Physiol 298: R862–869.

Tacconelli S, Patrignani P (2014) Inside epoxyeicosatrienoic acids and cardiovascular disease. Front Pharmacol 5: 239.

Tacke F, Zimmermann HW (2014) Macrophage heterogeneity in liver injury and fibrosis. J Hepatol 60: 1090–1096.

Tak PP, Firestein GS (2001) NF-kappaB: a key role in inflammatory diseases. J Clin Invest 107: 7–11.

Tan H, He Q, Li R, Lei F, Lei X (2016) Trillin reduces liver chronic inflammation and fibrosis in carbon tetrachloride (CCl_4_) induced liver injury in mice. Immunol Invest 45: 371–382.

Taniguchi K, Karin M (2018) NF-kappa B, inflammation, immunity and cancer: coming of age. Nat Rev Immunol 18: 309–324.

Theken KN, Deng Y, Kannon MA, Miller TM, Poloyac SM, Lee CR (2011) Activation of the acute inflammatory response alters cytochrome P450 expression and eicosanoid metabolism. Drug Metab Dispos 39: 22–29.

Tsunedomi R, Iizuka N, Hamamoto Y, Uchimura S, Miyamoto T, Tamesa T, Okada T, Takemoto N, Takashima M, Sakamoto K et al (2005) Patterns of expression of cytochrome P450 genes in progression of hepatitis C virus-associated hepatocellular carcinoma. Int J Oncol 27: 661–667.

Uhlén M, Fagerberg L, Hallström BM, Lindskog C, Oksvold P, Mardinoglu A, Sivertsson Å, Kampf C, Sjöstedt E, Asplund A et al (2015) Proteomics. Tissue-based map of the human proteome. Science 347: 1260419.

Wagner EF, Nebreda AR (2009) Signal integration by JNK and p38 MAPK pathways in cancer development. Nat Rev Cancer 9: 537–549.

Wang X, Yu T, Liao X, Yang C, Han C, Zhu G, Huang K, Yu L, Qin W, Su H et al (2018) The prognostic value of CYP2C subfamily genes in hepatocellular carcinoma. Cancer Med 7: 966–980.

Xia J, Sinelnikov IV, Han B, Wishart DS (2015) MetaboAnalyst 3.0--making metabolomics more meaningful. Nucleic Acids Res 43: w251–257.

Xu XR, Huang J, Xu ZG, Qian BZ, Zhu ZD, Yan Q, Cai T, Zhang X, Xiao HS, Qu J et al (2001) Insight into hepatocellular carcinogenesis at transcriptome level by comparing gene expression profiles of hepatocellular carcinoma with those of corresponding noncancerous liver. Proc Natl Acad Sci USA 98: 15089–15094.

Ye H, Liu X, Lv M, Wu Y, Kuang S, Gong J, Yuan P, Zhong Z, Li Q, Jia H et al (2012) MicroRNA and transcription factor co-regulatory network analysis reveals miR-19 inhibits CYLD in T-cell acute lymphoblastic leukemia. Nucleic Acids Res 40: 5201–5214.

Yu D, Green B, Marrone A, Guo Y, Kadlubar S, Lin D, Fuscoe J, Pogribny I, Ning B (2015) Suppression of CYP2C9 by microRNA hsa-miR-128-3p in human liver cells and association with hepatocellular carcinoma. Sci Rep 5: 8534.

Zhang HM, Kuang S, Xiong X, Gao T, Liu C, Guo AY (2015) Transcription factor and microRNA co-regulatory loops: important regulatory motifs in biological processes and diseases. Brief Bioinform 16: 45–58.

Zhou L, Du Y, Kong L, Zhang X, Chen Q (2018) Identification of molecular target genes and key pathways in hepatocellular carcinoma by bioinformatics analysis. Oncotargets Ther 11: 1861–1869.

